# Differences in binding preferences for XIST partners are observed in mammals with different early pregnancy morphologies

**DOI:** 10.1101/2023.10.04.560814

**Authors:** Ioannis Tsagakis, Haidee Tinning, Irene Malo-Estepa, Adrian Whitehouse, Mary J. O’Connell, Niamh Forde, Julie L Aspden

## Abstract

In comparison to protein-coding genes, long non-coding RNAs (lncRNAs) are poorly conserved between species at the sequence level yet they often perform conserved and essential regulatory roles. The lncRNA *XIST* mediates X chromosome inactivation (XCI) through the interaction of numerous proteins, most of which bind to its repeat regions. Whilst *XIST* is present across placental mammals, its sequence is divergent. In addition, the timing and mechanistic details of the process of XCI also vary across placental mammals. Here, we sought to determine whether *XIST* interactor proteins previously identified in a model mammalian species (mouse) also bind in other mammals which exhibit divergent timing of XCI (human and bovine) and differing pre-implantation embryo morphology. We show that *XIST* and its putative protein partners are coordinately expressed in the endometria of mammals with different pre-implantation embryo morphologies. RNA immunoprecipitations revealed that human WTAP, SPEN, hnRNPK and CIZ1 proteins bind human *XIST*. CIZ1-*XIST* binding was also detected in bovine. In both human and cow, we found CIZ1 binds to the repeat E element of *XIST*. However, human CIZ1 protein is not able to bind bovine repeat E, indicating a species-specific binding interaction. Together these data indicate that several of the key RNA-protein interactions with *XIST* are common across placental mammals, even though we observe divergence in both the RNA sequence of *XIST*, and early embryo morphologies. This work sheds light upon the evolution of RNA-protein binding interactions, revealing that the binding events can be conserved even when the precise mechanism of binding has changed.

**Statements and Declarations:** Authors have no financial or non-financial interests that are directly or indirectly related to the work submitted for publication.

## INTRODUCTION

Maintaining gene dosage across sexes in mammals is vital as an imbalance can result in embryonic lethality (Marahrens et al. 1997). This process of X-Chromosome Inactivation (XCI) in mammals is mediated via a long non-coding RNA (lncRNA) *X-inactive specific transcript* (*XIST),* the length of which varies across species e.g., from 17 to 35 kb lncRNA in humans and bovine respectively (Brockdorff 2018). The majority of what we know about XCI and *XIST* is from data in the mouse which is developmentally anomalous in mammals (Ramos-Ibeas et al. 2019). Across *XIST*’s first and last exons, repetitive regions serve as its ‘functional domains’ (Brockdorff 2018) with repeat regions A, B, C and E containing protein binding sites for up to 81 proteins partners, at least in mouse (Chu et al. 2015). Once *XIST* is expressed in a cell, it recruits a multitude of protein partners to orchestrate the repression of active genes on the X chromosome to achieve dosage compensation (Jégu, Aeby, and Lee 2017). This is essential to equalise the expression of genes on the X chromosome between males and females. Gene dosage imbalance is deleterious for embryo development (Takagi and Abe 1990) and female mice without the Xist locus all die by ∼18 days (Yang et al. 2016).

Of the known protein partners for *XIST* (Figure 1A), Spen interacts with the A-repeat in mouse embryonic stem cells (Chen et al. 2016; Lu et al. 2016; Lu et al. 2020; Monfort et al. 2015) and in human embryonic kidney cells (HEK293T) (Graindorge et al. 2019) mediating the silencing of active genes on an X chromosome. RBM15 and WTAP proteins also interact with the A-repeat of mouse *Xist* (Chu, et al. 2015; McHugh et al. 2015), mediating X-linked gene silencing via recruitment of the m6A methylation machinery (METTL3/4) to *XIST* in HEK293T cells (Patil et al. 2016). In contrast, hnRNPK binds repeat B and C of mouse *Xist* (Almeida et al. 2017; Pintacuda et al. 2017; Bousard et al. 2019) and human hnRNPK interacts with human *XIST* B, C and D repeats (Lu, et al. 2020). hnRNPK links PRC1/PRC2-mediated repressive chromatin modification (H2AK119ub1 and H3K27me2/3) installation on active X-linked genes. HnRNPU binds *Xist* in mouse Neuro2A (neural crest-derived) cells (Hasegawa et al. 2010) and HEK293T cells (Lu, et al. 2020), contributing to XCI by localising *Xist* on the X chromosome. Mouse Ciz1 binds *Xist* repeat E enabling anchoring of *Xist* to the nuclear matrix (nuclear periphery) and to the inactive X chromosome (Xi), facilitating gene silencing (Ridings-Figueroa et al. 2017; Sunwoo et al. 2017). Lbr was also identified as a bona fide interacting partner of mouse *Xist*, interacting via its repeat A region (McHugh, et al. 2015; Lu, et al. 2020) and tethers *Xist* in the cell’s nuclear lamina (Chen, et al. 2016). Therefore, it is clear that effector proteins are necessary for *Xist* to initiate and establish XCI in mouse. However, details of which protein partners are involved in XCI in other mammals is currently lacking and for human *XIST* interactions described have only been characterised in less relevant cell lines (HEK293, K562, HepG2, GM12878 cells) (Figure 1B).

**Figure 1.**
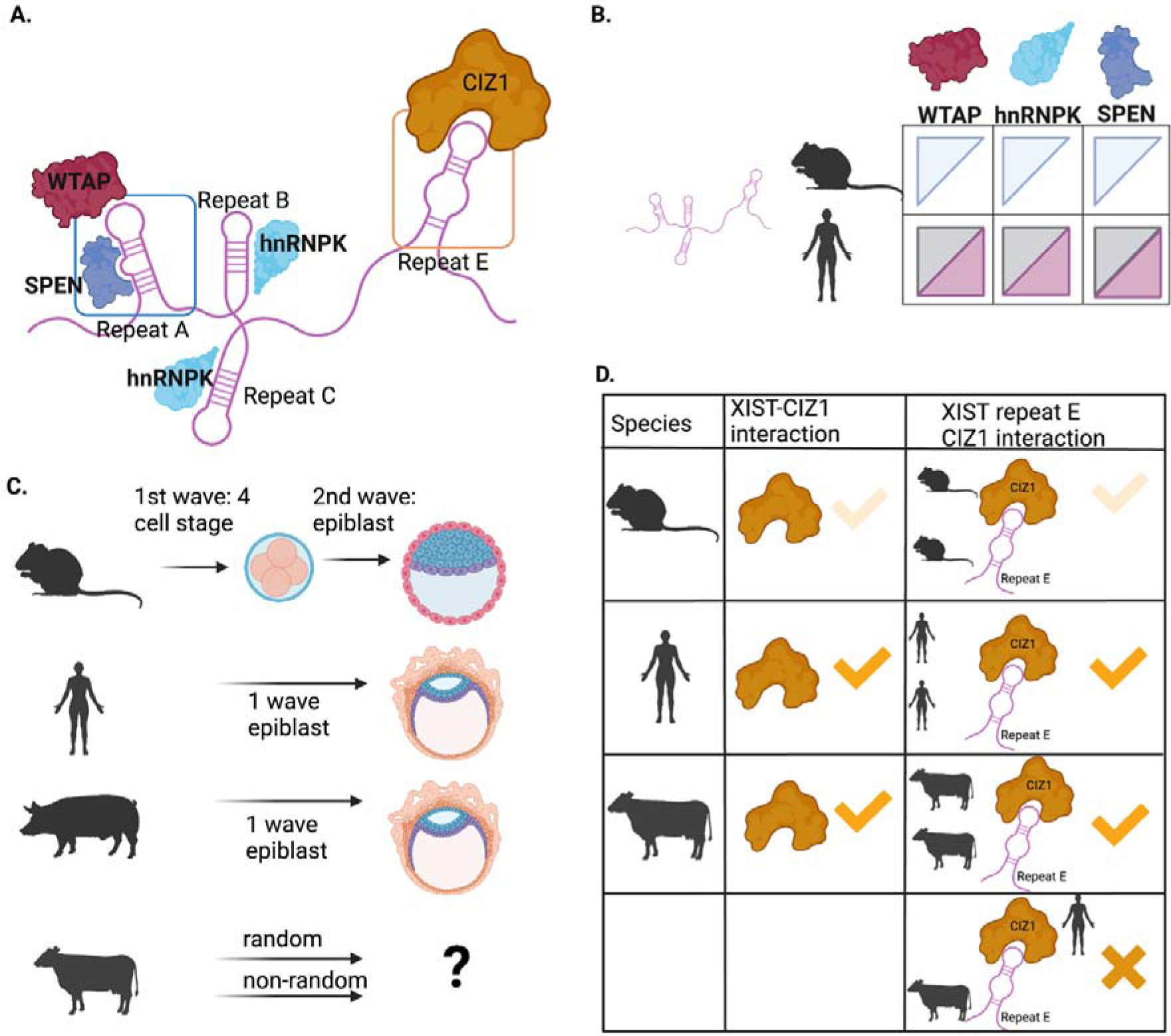
Overview of *XIST*-protein interactions and conservation across different mammals. A) Schematic of key *XIST*-protein interactions investigated here including sites of characterised interactions based on work in mouse and human HEK293 cells. B) Summary of evidence for specific protein-*XIST* interactions in mouse ESC (blue triangles), and human HEK293 cells (grey triangles), along with summary of interactions established here in human Ishikawa cells (pink triangles). C) Overview of different embryonic and pregnancy morphologies. D) Summary of characterisation of *XIST*-CIZ1 interactions from literature (mouse in light orange) and our results in human Ishikawa cells and bovine stromal cells (dark orange), as well as cross species testing. Created with BioRender.com

LncRNAs have diverse functions depending on where the lncRNA is localised and also on what molecule they interact with, including protein partners (Tsagakis et al. 2020). Contrary to most lncRNAs, which exhibit rapid evolution and thus low levels of sequence conservation (Pang, Frith, and Mattick 2006), *XIST* has moderate conservation among the eutherian clade (Yen et al. 2007). Yet our current knowledge regarding the role of *XIST* and its interaction partners in XCI comes from either *in vivo* mouse models, mouse or human stem cells or generic human tissue culture cell lines (e.g. HEK293), which are unlikely to be representative of the entire eutherian clade. For instance, certain XCI milestones such as temporal *XIST* expression and X-linked gene silencing are achieved at different time-points in different placental mammal species (Okamoto et al. 2011; Vallot, Ouimette, and Rougeulle 2016) (Figure 1C). The timing of implantation differs among placental mammals with the mouse and human embryo implanting in the uterus five and seven days after fertilisation respectively, whereas pigs and cattle could take up to two to three weeks (Berg et al. 2010; Bou et al. 2017). Furthermore, the nature of XCI varies (imprinted vs random) along with the extent of XCI escape across placental mammals. In mice, there is a wave of imprinted XCI where the paternal X is preferentially briefly inactivated from the 4-cell stage to the blastocyst until the paternal X is reactivated transiently in preparation for the second wave of random XCI (reviewed in (Vallot, Ouimette, and Rougeulle 2016)). In contrast, humans (Petropoulos et al. 2016) and pigs (Zou et al. 2019) experience a single wave of random XCI whereas the picture is unclear in cow (Chen, et al. 2016; Xue et al. 2002; Dindot et al. 2004; Couldrey et al. 2017; Figure 1C).

A statement of sequence homology between genes describes shared ancestry but does not directly transfer functional information. Even single gene orthologs can have divergent functions through time and in different lineages (e.g. (Gharib and Robinson-Rechavi 2011)). By comparing alignments of homologous sequences across species we can gain insight into variation in selective pressure acting in site and/or lineage specific ways which can underpin species/lineage-specific divergence in function of the corresponding protein products. The vast majority of protein coding regions evolve under a strict selective constraint to maintain organismal fitness where substitutions that emerge are actively purged by natural selection (*i.e.* purifying or negative selection). On the contrary, substitutions that increase fitness are “positively selected” and increase in frequency in the population through time. In general, observations of positive selection are considered synonymous with functional shift (Levasseur et al. 2006). Rational mutagenesis experiments and functional assays designed to assess the impact of residues under positive selection support this thesis. For example, the vertebrate umami taste receptor has been co-opted in hummingbirds to function as a sweet taste receptor, and in mammals the evolution of the unique chlorination activity of myeloperoxidase (MPO) (an innate immune enzyme produced by neutrophils) (Baldwin et al. 2014; Loughran et al. 2012). By assessing the selective pressure variation across species for a given set of orthologs we can therefore predict whether there is evidence for positive selection and hence protein functional shift. Here we test the hypothesis that homologs of protein interacting partners of *XIST* are under the similar selective constraints across species with different morphology and timing of XCI.

Therefore, under the prism that eutherian mammals have diverged in terms of their early development, XCI timing, and reproductive morphologies, it is possible that their mechanisms of XCI have diverged as well, perhaps co-opting new protein partners. We tested the hypothesis that the protein partners that *XIST* interacts with, are different in species with different morphology and timing of XCI. Given XCI only occurs in females and the expression of *XIST* is higher in female reproductive tissues we selected to dissect the *XIST*-protein interactions in a human endometrial adenocarcinoma cell line (Ishikawa) and bovine stromal cells from the endometrium. Our aim was to characterise *XIST*-protein interactions within the context of the human and cow endometrium and to provide insight into XCI within the context of reproductive tissue and placental mammals other than mouse (Figure 1D).

## MATERIALS AND METHODS

Unless stated otherwise, all consumables were sourced from Sigma-Aldrich.

### Analysis of sequence conservation of XIST and its protein partners

We sought to determine how conserved the sequences of *XIST* and its protein partners were in selected mammalian species with different developmental morphologies. The RNA sequences of *XIST* and its mouse protein partners were sourced from Ensemble 90 or later. Where sequences were not found in Ensemble, sequences from GenBank were used. More specifically, the following RNA sequences were used for *XIST*: human (NR_001564.2), mouse variant 1 (NR_001463.3), pig (KC753465.1) and predicted bovine variant 1 (XR_001495594.1). The following peptide sequences were retrieved from UniProt for mouse, human, pig and cow, respectively: SPEN (Q96T58, Q62504, A0A287BPC2, F1MRK2), WTAP (Q15007, Q9ER69, F1SB61, F1MN80), RBM15 (Q96T37, Q6PHZ5, F1S619, E1BFX6) and CIZ1 (Q9ULV3, Q8VEH2, F1RRW9, F1MZB8).

To compare the sequence identities of *XIST* RNA sequences across the four species and the mRNA sequences coding for protein partners of mouse *Xist*, the multiple sequence alignment software Clustal Omega (https://www.ebi.ac.uk/Tools/msa/clustalo/) was used with the default settings to derive percent identity matrix scores. Alignments of peptides and their domains were performed with Clustal-ω with default alignment, numbering and colour options.

### Endometrial cells/tissue isolation

To determine if *XIST* and its protein partners were expressed coordinately in endometrial tissue we generated samples from four species 1) human, 2) cow, 3) pig, and 4) mouse. Human immortalised endometrial epithelial cells (Ishikawa cells) were cultured at 37°C (5% CO_2_) in DMEM:F12 B-Nut mix (50:50, ThermoFisher) supplemented with 10% (v/v) heat-inactivated Fetal Bovine Serum and 5% (v/v) Penicillin-Streptomycin-Glutamine. Cow endometrial tissue was isolated from the ipsilateral uterine horn at the late luteal stage of the estrous cycle from female *Bos taurus* (local abattoir), and cow endometrial stromal cells were isolated and maintained as described by (Tinning et al. 2020). Pig endometrial tissue was isolated from uteri of females (n=3) slaughtered at bacon weight from a local abattoir. Mouse uteri (n=3) were collected from unknown age female NFAT-Luc ApoE *Mus musculus* following a schedule one procedure. All tissues were snap-frozen in LN_2_ and stored at -80°C. For cytoplasmic/nuclear fractionation, ∼30 million Ishikawa cells were lysed in a hypotonic lysis buffer (10 mM HEPES [ThermoFisher], 1.5 mM MgCl_2_, 1 mM KCI, 0.5 mM DTT, supplemented with 1X EDTA-free protease inhibitor cocktail] for 40 minutes on ice. Lysates were centrifuged at 800 xg for 8 minutes and cytoplasmic supernatant (cytoplasm) transferred into fresh tubes and centrifuged twice more whereas the nuclear pellet was resuspended in 1 ml of ice-cold hypotonic lysis buffer. The resuspended pellet was kept on ice for 5 minutes before centrifuging at 800 xg for 8 minutes and then resuspending in 500 μl of 1X PBS. Endometrial tissue pieces (from cow, pig, and mouse) were homogenised via a mechanical rotor (Heidolph SilentCrusher S) in 600 µl of RNA lysis buffer (mirVana miRNA isolation) and total RNA was isolated using the same kit, according to the manufacturer’s instructions.

### RT-qPCR expression profiling of XIST and putative protein partners

One µg of RNA per sample was used to generate cDNA using the High-Capacity cDNA Reverse Transcription, (ThermoFisher), according to the manufacturer’s instructions, in a Veriti PCR thermocycler (Applied biosystems). Samples were cycled once at 25°C for 10 minutes, 37°C for 2 hours and 85°C for 5 minutes. Gene expression quantification proceeded with a SYBR Green Mastermix (Roche Diagnostics LightCycler 480 SYBR Green I) and LightCycler 480 II (Roche). A total of 12.5 ng of cDNA were added per well and serial dilutions of 5 were prepared from a 1:10 diluted pool of cDNA derived from all cDNA samples assayed. Final primer concentrations were 0.5 μM (Supplementary Table S1). Cycling conditions were: 95 for 5 minutes (x1), 95°C for 10 seconds, 60°C for 10 seconds and 72°C for 10 seconds (x45 cycles). Melt curves were estimated by heating for 5 seconds at 95°C, 65°C for 1 minute followed by a continuous increase to 97°C while acquiring fluorescence readings (5/°C). Expression levels obtained for the various genes of interest assayed in the different systems were normalised to β-actin (*Actb*) from each species.

### Protein lysate generation for immunoblotting profiling of putative XIST protein partners

To generate protein lysates, 10 million Ishikawa cells were lysed in 250 μl of RIPA buffer (150 mM NaCl, 1% IGEPAL, 0.5% Sodium deoxycholate, 0.1% SDS, 25 mM Tris-HCl [pH 7.4], supplemented with protease inhibitor cocktail in water). Cell lysates were incubated on ice for 10 minutes and passed five times through a 21’ G needle and syringe to homogenise. Cell lysates were centrifuged at max speed (17,000 xg) for 1 minute to remove insoluble material and supernatants were frozen at -20°C. All endometrial tissue pieces were homogenised via a mechanical rotor (Heidolph SilentCrusher S) in 600 µl of RIPA buffer and protein lysates were generated as described above. Protein concentration was quantified using the Pierce™ BCA Protein Assay Kit (according to the manufacturer’s instructions) and absorbance measured on a plate reader at 595 nm. Protein samples were diluted with 6x Laemmli buffer and heated at 95°C for 5 minutes, separated on a 10% denaturing SDS-polyacrylamide gel at 90 V for 30 minutes and 150 V for 80 minutes using 1x running buffer (1x running buffer, 0.025 M Tris, 0.25 M Glycine and 0.1% SDS in water). Subsequently, proteins were wet-transferred onto a 0.22 μm PVDF membrane at 200 mA for 1:30 hours using 1x transfer buffer (25 mM Tris and 192 mM Glycine with 20% methanol in water). Membranes were blocked with 5% Marvel milk powder in PBS-T (0.5% Tween) for 1 hour at room temperature while rocking. Blots were incubated with primary antibodies (Supplementary Table S2) overnight at 4°C while rolling, washed three times in PBS-T (0.5% Tween) at room temperature for 10 minutes each while rolling and incubated with secondary HRP-conjugated antibodies for 1 hour at room temperature while rolling. Membranes were washed three times in PBS-T (0.5% Tween) at room temperature for 10 minutes each while rolling and developed using ECL.

#### RIP coupled to RT-qPCR in human and cow

Cell pellets of 40 million Ishikawa or bovine stromal cells were snap-frozen and stored at -80°C. Cells were lysed with 100 μl with RIP lysis buffer (Magna RIP kit, Merck) supplemented with 10 U RNaseIn and 0.5 μl protease-inhibitor cocktail; per 20 million cells while incubating on ice for 15 minutes, with frequent inversion. Lysates were then stored at -80°C. Lysates were passed through a 27’ gauge needle and syringe 3-5 times to homogenise. Pooled lysates were clarified at 17,000 xg for 1 minute. 50 μl of magnetic bead suspension were washed twice with 500 μl of RIP Wash Buffer. Beads were resuspended in 100 μl RIP Wash Buffer and mixed with ∼5 μg of antibody (apart from ∼1.2 μg for hnRNPK). Antibody-bead mixtures were incubated for 30 minutes at room temperature while rotating before washing three times with RIP Wash Buffer A.

For each RIP reaction, 900 μL of RIP Immunoprecipitation Buffer (Magna RIP kit, Merck) supplemented with 200 U RNaseIn was mixed with 100 μl of the lysate and this mixture was used to resuspend the antibody-coupled beads. Reactions were incubated overnight at 4°C while rotating. Supernatant was kept for RNA and protein downstream analyses (“depleted lysate”) and RNP complexes on beads were washed with 500 μl of RIP Wash Buffer six times. 10% of beads were resuspended with 1x of Laemmli buffer and analysed by western blot. The equivalent of 500,000 cells were used for input and depleted lysate as well as whole elution. To elute RNA, beads were treated with 150 μl of Proteinase K buffer, incubated at 55°C for 30 minutes while shaking at 1100 RPM and supernatant was kept for RNA analyses.

250 μl of ddH2O was added to eluted RNA and mixed with 400 μl of phenol:chloroform:isoamyl alcohol while vortexing for 15 seconds. Samples were centrifuged at 17,000 xg for 10 minutes at room temperature. The aqueous layer was retrieved and mixed with 400 μl of chloroform. Samples were vortexed for 15 seconds and centrifuged at 17,000 xg for 10 minutes at room temperature, after which the aqueous layer was mixed with 920 μl of Ethanol Precipitation buffer (supplemented with 2 μl of GlycoBlue; ThermoFisher) and precipitated at -80°C for >3 hours. Samples were centrifuged at 17,000 xg for 30 minutes at 4°C and RNA pellets washed with 1 ml of 75% ethanol. RNA pellets were resuspended in 15 μl of RNase-free water and quantified by NanoDrop1000 instrument. 500 ng of RNA from Ishikawa or bovine stromal cells were used per sample to make cDNA (for SPEN, 1000 ng were used per sample). Primers were designed for *XIST* and control transcripts for human and bovine (Supplementary Table S3). Fold enrichment of each transcript’s abundance in RIP elutions was normalised to input using the 2^^-(Ct^_elution_^- Ct^_input_^)^ formula and reported as RNA enrichment.

#### Cloning of human and bovine *XIST* repeats

Human and bovine *XIST* E repeats were PCR amplified from cDNA generated from either total RNA of Ishikawa (human) or bovine stromal cells (Supplementary Table S4) with the Q5 High-Fidelity DNA polymerase (NEB, UK) or the non-proofreading Taq DNA Polymerase (EP040; ThermoFisher, UK) and inserted into TOPO Blunt vector (Zero Blunt™ TOPO™ PCR Cloning Kit; ThermoFisher, UK), downstream of an SP6 promoter and upstream of a T7 promoter. Plasmid sequences were confirmed using Sanger sequencing.

#### *In vitro* transcription of human and bovine *XIST* repeats biotinylated nucleotides

Plasmids were linearised with either EcoRV or BamHI to generate sense and anti-sense RNAs (Supplementary Table S5). RNA was transcribed using 1 μg of linearised plasmid DNA, 1x of Biotin RNA labelling mix (1x mixture: 1 mM ATP, 1 mM CTP, 1 mM GTP, 0.65 mM UTP, 0.35 mM Biotin-16-UTP, pH 7.5; Roche/Sigma) and ∼40 U of either T7 (Roche/Sigma, UK) or SP6 (Roche/Sigma, UK) polymerase in 20 μl total volume at 37°C for 2 hours. For the SP6 polymerase, the HiScribe SP6 RNA Synthesis (NEB, UK) kit was also used. Reactions were -treated using 6 U of Turbo DNase (ThermoFisher, UK) at 37°C for 15 minutes. RNA was purified using chloroform and isopropanol precipitated. Transcript sizes were checked on 1% denaturing formaldehyde agarose gels in 1xMOPS buffer (20 mM MOPS pH 7.0, 12 mM sodium acetate, 0.5 mM EDTA pH 8.0) alongside RNA ladder (RiboRuler High Range; NEB, UK). Biotinylation was confirmed via RNA slot blot.

#### Ishikawa nuclear extract

10 million Ishikawa were fractionated as previously with a few modifications (Werner and Ruthenburg, 2015). Briefly, cell pellets were resuspended in 250 μl of Buffer A (10 mM HEPES-KOH pH 7.5, 10 mM KCl, 10% glycerol, 340 mM sucrose, 4 mM MgCl_2_, 1 mM DTT in ddH2O supplemented with 1x cOmplete™, Mini, EDTA-free protease inhibitor cocktail; Sigma, UK) and 250 μl of Buffer A+Triton (2% Triton-X100, 10 mM HEPES-KOH pH 7.5, 10 mM KCl, 10% glycerol, 340 mM sucrose, 4 mM MgCl_2_, 1 mM DTT in ddH2O supplemented with 1x cOmplete™, Mini, EDTA-free protease inhibitor cocktail). The cell suspension was incubated on ice for 12 minutes with occasional inverting to mix followed by centrifugation at 1,200 xg for 5 minutes at 4°C. Nuclear pellets were resuspended in 250 μl of Buffer A and 250 μl of Buffer A+Triton and centrifuged at 1,200 xg for 5 minutes at 4°C. Nuclear pellets were resuspended with 250 μl of RNP lysis buffer (20 mM Tris-HCl pH 7.5, 500 mM LiCl, 0.5% LiDS, 1 mM EDTA, 5 mM DTT in ddH2O, supplemented with protease inhibitor cocktail and 1U/μl RNase inhibitor) and incubated on ice for 20 minutes. Nuclear extracts were then homogenised by passing through a 27’ gauge needle and syringe 5-7 times and then centrifuged at 17,000 xg for 1 minute. Protein concentration was determined via the Protein Qubit assay.

#### *In vitro* transcribed RNA pulldown

Nuclear-enriched extracts from Ishikawa or whole-cell extracts from bovine stromal cells were used. Each lysate containing 0.5-1 mg protein was mixed with the incubation buffer (20 mM Tris-HCl pH 7.5, 150 mM NaCl, 1.5 mM MgCl_2_, 2 mM DTT, 0.5% sodium deoxycholate, 0.5% NP-40 in ddH2O supplemented with protease inhibitors and 1 U/μl RNase Inhibitor). The ratio of lysis buffer to incubation buffer was ∼1:4. Lysates were pre-cleared by incubating with 50 μl of Dynabeads™ MyOne™ Streptavidin C1 magnetic beads (10 mg/ml; ThermoFisher, UK). 10 μg of biotinylated RNA was denatured at 85°C for 3 minutes and snap-chilled on ice for 2 minutes before incubating with pre-cleared lysates for 2 hours while rotating at 4°C. 50 μl of beads were then added and incubated for further 1 hour at 4°C while rotating to recover biotinylated *XIST*-protein complexes. Beads were washed six times with washing buffer (20 mM Tris-HCl pH 7.5, 150 mM NaCl, 1.5 mM MgCl_2_, 2 mM DTT, 0.5% sodium deoxycholate, 0.5% NP-40 in ddH2O). For downstream analyses via western blot, beads were resuspended in 1x Laemmli buffer and heated at 95°C for 5 minutes while shaking at 1100 RPM.

### Comparative Genomics of *XIST* protein interacting partners

#### Identification of homologs of interacting protein partners across mammals

Placental mammal genomes included in this work were chosen based on the quality of the genome available and their phylogenetic placement. In total 16 placental mammal species genomes were used (Macaque, Gorilla, Orangutan, Chimpanzee, Human, Mouse, Rabbit, Microbat, Horse, Pig, Cat, Dog, Dolphin, Cow, Elephant, and Armadillo), one marsupial (Opossum), and one monotreme (Platypus) (Supplementary Table S6). The 9 protein-coding genes selected here were SPEN, WTAP, RBM15, CIZ1, hnRNPK, hnRNPU, PTBP1, MATR3 and LBR. The OMA database (Altenhoff et al. 2018) was queried for 1:1 orthology and the corresponding Coding DNA sequences (CDSs) for single gene orthologs were retrieved from Ensembl 102 (Yates et al. 2020). The corresponding protein coding sequences were generated using MAFFT (Katoh and Standley 2013).

#### Analysis of lineage-specific selective pressure variation

Using the VESPA pipeline (Webb, Walsh, and O’Connell 2017) files were generated in the appropriate format to carry out selective pressure variation analysis. Variation in selective pressure was assessed using codon-based models of evolution to identify changes in dN/dS across sites and lineages as implemented in codeml in the PAML package (v4) (Yang 2007). The models we employed are a set of standard nested models which are automatically compared by Vespa using likelihood ratio tests with significance calculated using the appropriate degrees of freedom. More specifically, the models used were the neutral model M1Neutral, and its lineage-specific extensions model A, and the null model for model A. M1Neutral allows two site classes for dN/dS (referred to as ω throughout): ω0=0 and ω1=1. Model A assumes the two site classes are the same in both foreground and background lineages (ω0=0 and ω1=1) and ω2 for the foreground is estimated from the data and free to vary above 1. Model A null estimates a ω2 value also, but here it is restricted to below 1 thus allowing sites to be evolving under either purifying selection, or to be neutrally evolving but not permitting positive selection. Sequences were considered to exhibit lineage-specific selective pressure if the likelihood ratio test (LRT) for ModelA is significant in comparison to both ModelA null and M1Neutral. The lineages labelled as foreground were human, mouse, cow and pig.

In all cases where models allowed for the estimation of positively selected sites, the contributions of individual sites to the signal for positive selection was estimated using a Bayes Empirical Bayes (BEB) approach. BEB estimates for each site on each gene were reported if the posterior probability was greater than 0.5. To obtain a set of identified sites with higher confidence, two user-defined posterior probability (PP) cut-offs were set at PP=0.95 and PP=0.99.

## RESULTS

### Coordinate expression of XIST and its protein partners in the endometrium

To examine how the sequence of *XIST* and its putative protein partners varies across placental mammals, we employed Clustal ω (Sievers et al. 2011) to determine the conservation of the *XIST* RNA sequence in the four species tested and estimated to be 63-73% (Figure 2A). Recent data indicate full-length lncRNA sequence conservation is not very informative for lncRNAs, as conservation of smaller regions across the transcript can predict function more reliably (Kirk et al. 2018). Aligning the sequences from *XIST*’s repeat regions confirmed that repeat A region displayed the highest % identity score ranging from ∼76% to 90% between the four species tested (Figure 2A). Repeat region B (62 to 91% depending on the species), and repeat region D were also highly conserved (67 to 84% across human, mouse, cow and pig; Figure 2A). Not all repetitive regions found in human or mouse have been identifed in cow or pig. However, the C, E, and F repeat regions are moderately conserved in the species they have been mapped to (Figure 2A). To quantify the expression levels of *XIST* in uterine tissue, data from the GTEx portal (https://www.gtexportal.org/home/), which contains tissue-specific gene expression data from 54 non-diseased tissue sites across nearly 1000 human individuals were analysed. *XIST* abundance was highest in human reproductive tissues, including the endometrium and lowly expressed in other tissues, e.g. liver or blood (Figure 2B). Normalised expression levels of *XIST* were detected at low levels in human endometrial cells, and higher levels in mouse uterus, and consistently high levels in cow and pig endometrium (Figure 2C and Supplementary S7A).

**Figure 2.**
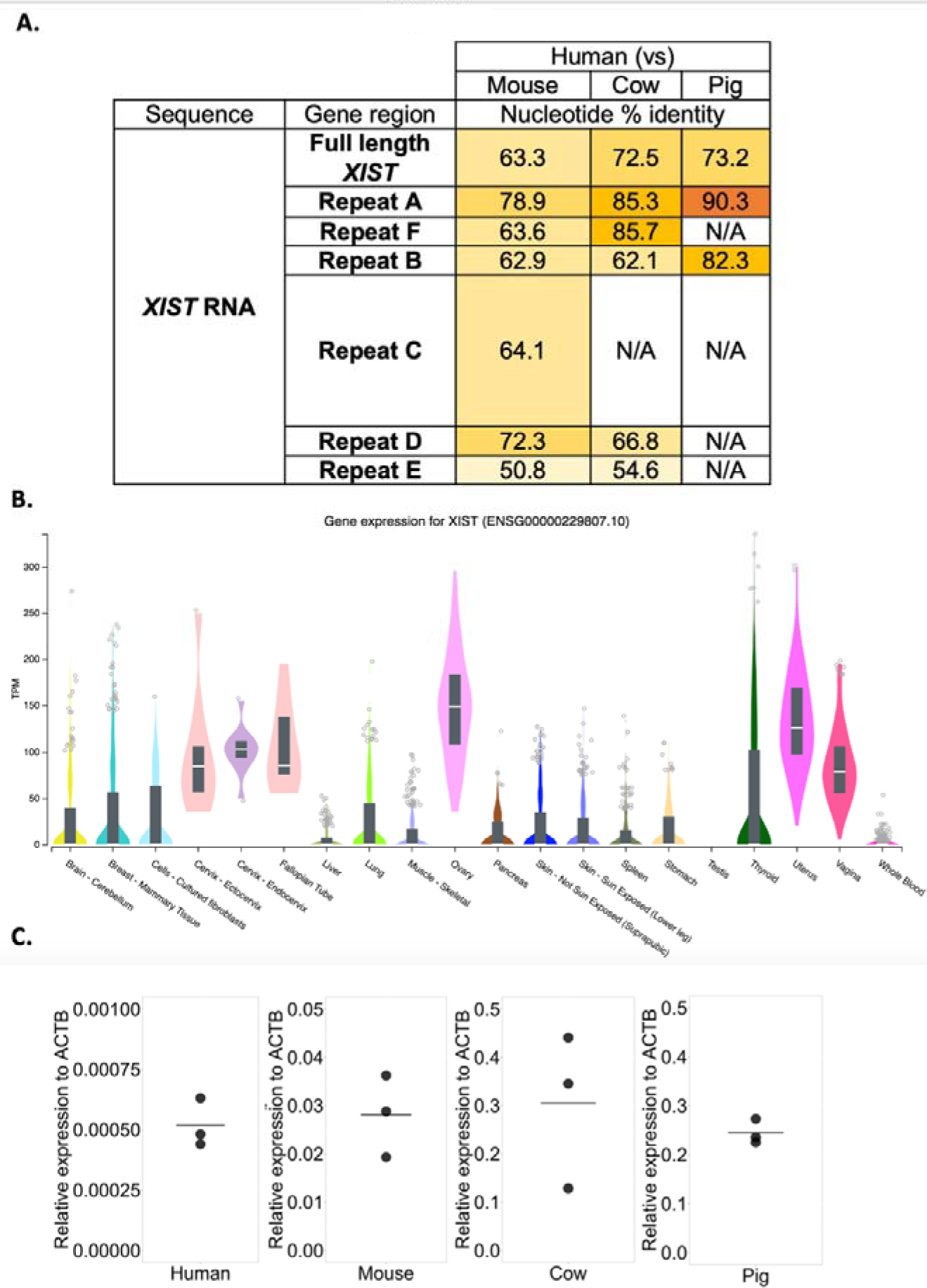
*XIST* sequence similarity and expression in selected mammals. **A)** Percent identity matrices aligning full-length *XIST* nucleotide sequences or repetitive regions on the *XIST* RNA from human, mouse, cow and pig with Clustal-ω (default settings). Percent identity scores are a proxy for conservation where a high % value shown represents similarity between the two compared sequences. N/A indicates that the repeat region is either not present or has not yet been mapped on the *XIST* sequence. Sequences were retrieved from Ensemble 90, or GenBank. Gene identifiers used: Mouse (NR_001463.3; GRCm38.p6); Human (NR_001564.2; GRCh38.p13); Cow (XR_001495594.1; ARS-UCD1.2); Pig (KC753465.1; Sscrofa11.1). **B)** Tissue-wide global RNA sequencing highlights *XIST (ENSG00000229807.10)* as most enriched in human reproductive tissues. Data obtained from https://gtexportal.org/home/gene/XIST on 29/12/2020. Expression values shown as Transcripts Per Million (TPM), calculated from a gene model with isoforms collapsed to a single gene. Box plots are shown as median and 25^th^ and 75^th^ percentiles. Outliers are displayed as dots if they are above or below 1.5 times the interquartile range. n=∼1000 human individuals. Colours indicate different tissue types and shades of same colour indicate tissue subtypes. **C)** *XIST* RNA levels measured by RT-qPCR (n=3 biological replicates per species) with species-specific *ACTB* as a reference gene in human Ishikawa cells, mouse uterus, cow endometrium and pig endometrium.

We also examined sequence similarity of protein partners previously identified to interact with *XIST* in mouse, specifically, Spen, Ciz1, Hnrnpk, Rbm15, Wtap, Lbr and Hnrpu proteins (Chu, et al. 2015; Hasegawa, et al. 2010; McHugh, et al. 2015; Ridings-Figueroa, et al. 2017; Sunwoo, et al. 2017; Monfort, et al. 2015; Kolpa, Fackelmayer, and Lawrence 2016; Pintacuda, et al. 2017; Moindrot et al. 2015; Yamada et al. 2015). At the amino acid sequence level, all proteins exhibited high similarity (>70%) between human, mouse, cow and pig: CIZ1 (∼70%) < LBR (∼80%), SPEN (∼80%) < RBM15 (∼95%), WTAP (∼95%) < hnRNPK, hnRNPU (∼99%)(Figure 3A). RT-qPCR indicated that mRNA from all 7 candidate genes was detected across human, mouse, cow and pig (Figure 3B). *SPEN, CIZ1, RBM15, WTAP* and *LBR* fluctuated at similar, low levels relative to *ACTB* across the four species. A pattern of high *hnRNPK* and *hnRNPU* expression relative to *ACTB* was seen consistently across all four species. Only a subset of putative *XIST* protein partners could be assayed via western blotting due to limited antibody availability for LBR and SPEN. Overall, signal was detected for all assayed proteins in all four species (Figure 3C). In summary, co-ordinate expression of both *XIST* RNA and putative protein partners was detected in the same cells and tissues at the same time in all four species tested. This raises the possibility that *XIST* from other placental mammals may interact with a subset of the same protein partners as was seen for the mouse.

**Figure 3.**
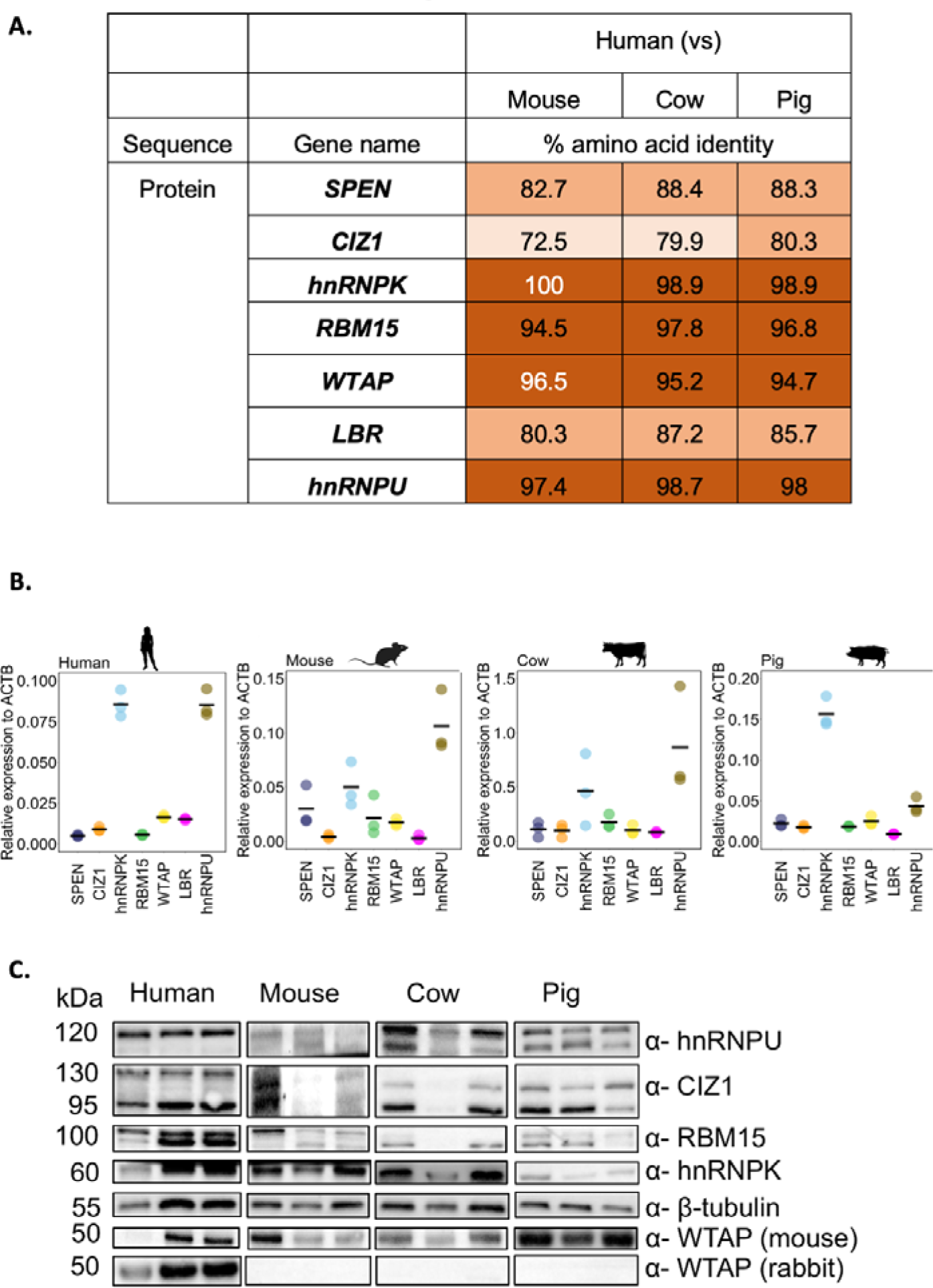
Sequence similarity and expression of protein partners known to interact with *XIST*. **A)** Percent identity matrices generated from amino acid sequences aligned with Clustal-ω (default settings) as a proxy for conservation and represent similarity between the two compared sequences. The default transition matrix was Gonnet, gap opening penalty = 6, gap extension = 1. Values in white text are candidates with higher levels of identity between human and mouse than between human and cow or pig. The darker the orange, the higher the level of amino acid identity. **B)** Relative mRNA expression levels of *XIST* protein partners measured by RT-qPCR with species-specific *ACTB* as a reference gene (n=3 biological replicates) in human Ishikawa cells, mouse uterus, cow endometrium and pig endometrium. **C)** Western blot analysis for *XIST* protein partners are present in cells and tissues of reproductive origin from human, mouse, cow and pig (n=3 biological replicates shown per species). Equal concentrations of protein (30 μg) were loaded per well with β-tubulin was used as a loading control. Blots containing three biological replicates from each species were probed separately for each protein per species as indicated by black boxes around blots.

### Human *XIST* interacts with human WTAP, hnRNPK and SPEN proteins

To determine whether the interactions identified between *XIST* RNA and protein partners in mouse occur in human cells, we performed RNA immunoprecipitations (RIP) followed by qRT-PCR. Specifically, we used antibodies against WTAP, hnRNPK and SPEN to pull these putative interactors down from human Ishikawa whole cell lysate and detect whether human *XIST* RNA was bound. These proteins were selected because of their functional contribution to XCI with *XIST* and their high levels of amino acid identity between mouse and human (Figure 3A).

The majority of WTAP was successfully immunoprecipitated from human Ishikawa whole cell lysate using two different WTAP antibodies (mouse and rabbit), with no non-specific binding to the IgG control. RT-qPCR primers were used to detect multiple exons of *XIST* (Supplementary 1A). RT-qPCR revealed that *XIST* was specifically associated with WTAP, compared to IgG control and negative control primers targeting *U2* and *ACTB* (Figure 4A). *XIST* RNA was also pulled down with hnRNPK, although *ACTB* control was also detected whilst U2 was not (Figure 4B). When compared to the IgG control this indicates that human *XIST* interacts with hnRNPK. Immunoprecipitation was also performed using a SPEN antibody, however the efficiency of pull-down could not be confirmed by western blotting because the antibody failed to recognise denatured protein. Therefore, mass spectrometry was used to determine if SPEN was precipitated. In the SPEN pulldown elution, 29 unique peptides were detected with a spectral count of 30, whilst no SPEN peptides were detected in the IgG control elution (Figure 4C). *XIST* RNA was found to be specifically bound to this immunoprecipitated SPEN compared to IgG control and negative control primer sets (Figure 4D).

**Figure 4:**
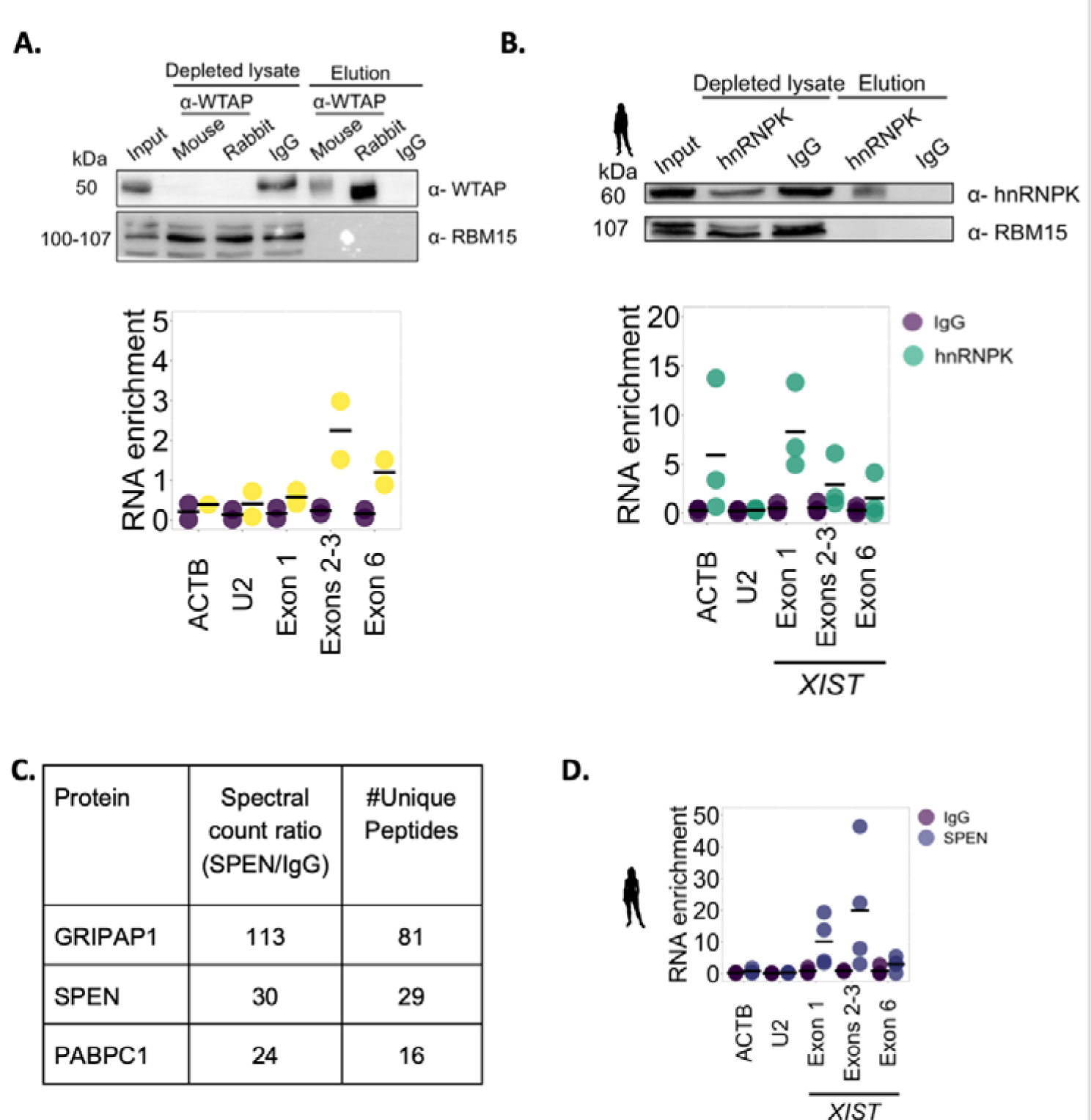
Human *XIST* interacts with WTAP, HNRNPK and SPEN. **A**) Representative image from western blot of WTAP RIP samples in whole cell extract of Ishikawa cells. Human-specific rabbit anti-WTAP antibody was used for western blotting and detects single WTAP isoform expected at ∼44 kDa and RBM15 used as negative control. Material loaded across input and depleted lysate samples was equivalent to ∼500,000 cells and 10% of the elution (n=2 biological replicates). Fold enrichment of transcript abundance in RIP elutions was normalised to input and reported as RNA enrichment. RT-qPCR results are from rabbit anti-WTAP antibody pull-down. Three technical replicates were performed for each of two biological replicates. IgG serves as a non-specific interacting protein negative control in RIP experiments. *ACTB* serves as a specific interacting transcript positive control whereas *U2* serves as non-specific transcript negative control. **B**) Representative western blot of hnRNPK RIP samples in whole cell extract of Ishikawa cells. One isoform expected at ∼60 kDa and RBM15 used as negative control. Material loaded across input and depleted lysate samples was equivalent to ∼500,000 cells and 10% of the elution (n=3 biological replicates). RT-qPCR fold enrichment of each transcript abundance in RIP elutions normalised to input and reported as RNA enrichment. Three technical replicates were performed for each of three biological replicates. IgG serves as a non-specific negative control in RIP experiments. *ACTB* serves as a specific interacting transcript positive control whereas *U2* serves as non-specific transcript negative control. **C**) Spectra count ratios for top 3 most abundant proteins in SPEN pull-down when compared to IgG control and number of unique peptides. SPEN is specifically pulled-down (n=1). **D**) RT-qPCR fold enrichment of each transcript abundance in SPEN RIP elutions normalised to input and reported as RNA enrichment. Three technical replicates were performed for each of four biological replicates. IgG serves as a non-specific negative control in RIP experiments. *ACTB* and *U2* serve as non-specific transcript negative controls.

RMB15 is a partner of *Xist* during XCI in mouse (binding through the A repeat). Although RBM15 protein was successfully immunoprecipitated from Ishkawa whole cell lysate there was little difference between *XIST*-RBM15 binding compared to IgG control (Supplementary S7B). To assess via an alternative method whether human *XIST* and RBM15 interact, biotinylated *XIST* RNAs were *in vitro* transcribed corresponding to either human *XIST* repeat A or an RNA corresponding to the anti-sense sequence of the repeat. These were then incubated in Ishikawa nuclear extracts and the ability of human RBM15 to interact with *XIST* repeat A was assessed by western blot compared with anti-sense negative control. Our results show that human RBM15 binds to *XIST* repeat A and not to the anti-sense sequence (Supplementary S7C). Together these results indicate that human *XIST* interacts with WTAP, hnRNPK, SPEN and RBM15, as previously characterised in mouse (Monfort, et al. 2015; Chu, et al. 2015; Almeida, et al. 2017).

### CIZ1 - *XIST* interaction via repeat E is conserved betwen human and bovine but species-specific in nature

To determine if human *XIST* interacts with CIZ1, as is reported in mouse, immunoprecipitations were performed with Ishikawa whole cell lysate. The majority of CIZ1 protein was successfully and specifically pulled down compared to IgG controls (Figure 5A). qRT-PCR at several sites across *XIST* showed it is specifically bound to CIZ1 compared to ACTB and U2 negative controls (Figure 5A).

**Figure 5:**
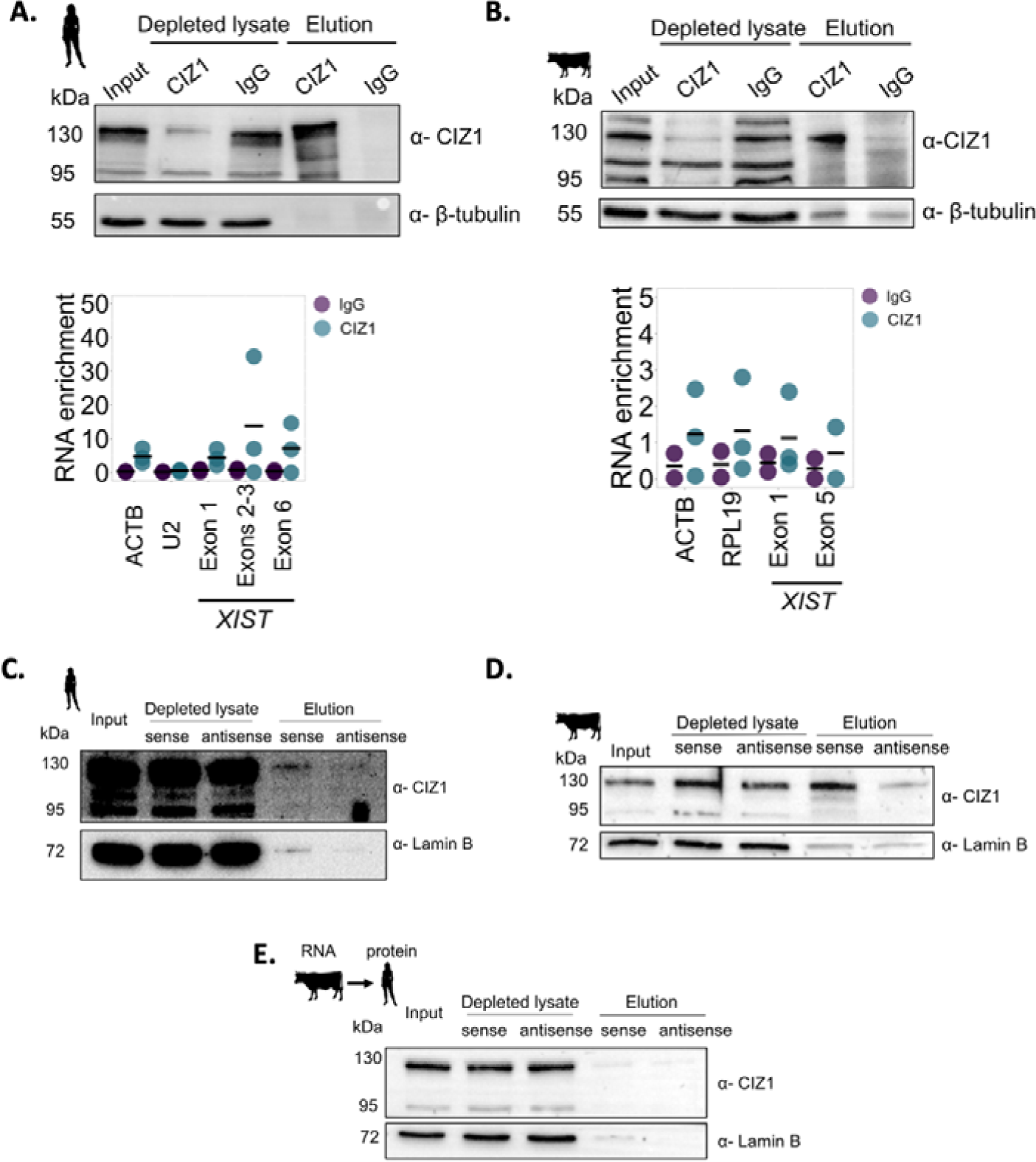
CIZ1 - *XIST* interaction via repeat E is conserved in human and bovine but species specific. **A**) Representative western blot of CIZ1 RIP from Ishikawa whole cell extract, with two CIZ1 isoforms successfully depleted (95kDa and 130 kDa). qRT-PCR revealed enrichment of *XIST* in CIZ1 pull-down compared to IgG compared to negative controls *ACTB* and *U2*. 3 technical replicates were performed for qRT-PCR from CIZ1 IP, on 3 biological replicates. **B)** Representative western blot of CIZ1 RIP from whole cell extract of bovine stromal cells, with two CIZ1 isoforms successfully depleted (95kDa and 130 kDa) (n=3 biological replicates). qRT-PCR revealed enrichment of *XIST* in CIZ1 pull-down compared to IgG compared to negative controls *ACTB* and *U2*. Three technical replicates were performed for each of three biological replicates. **C**) Representative western blot of *in vitro* transcribed biotinylated human *XIST* repeat E incubated with nuclear-enriched Ishikawa protein lysate. Equal amounts of protein were loaded from each sample (∼5 μg) and 100% of the elution sample. Input corresponds to 0.5%. Lamin B was used as a negative control, n=2 biological replicates. **D**) Representative western blot of *in vitro* transcribed biotinylated bovine *XIST* repeat E incubated with whole cell bovine stromal protein lysates. Equal amounts of protein were loaded from each sample (∼5 μg) and 100% of the elution sample. Input here corresponds to 0.5%. Lamin B was used as a negative control, n=3 biological replicates. Material from different animals was used for each biological replicate. **E**) Representative western blot of *in vitro* transcribed biotinylated bovine *XIST* repeat E incubated with nuclear-enriched Ishikawa protein lysate. Equal amounts of protein were loaded from each sample (∼5 μg) and 100% of the elution sample. Input here corresponds to 0.5%. Lamin B was used as a negative control, n= 2 independent biological replicates.

To establish if this CIZ1-*XIST* interaction occurs in other placental mammals, RIP qRT-PCR was also performed using bovine stromal whole cell lysate and the same CIZ1 antibody. Again the majority of CIZ1 protein was immunoprecipitated (Figure 5B). qRT-PCR across exons 1 and 5 of bovine *XIST* indicated that bovine *XIST* was enriched in CIZ1 pulldowns compared to IgG, although some signal was detected with negative control primers for *ACTB* and *RPL19* (Figure 5B). These results suggest that bovine *XIST* interacts with bovine CIZ1.

To identify the sites of these interactions on *XIST* we took an orthogonal approach to test whether it is the *XIST* repeat E element interacting with CIZ1 protein, as has been shown in mouse (Sunwoo, et al. 2017; Ridings-Figueroa, et al. 2017). Biotinylated *XIST* RNAs were *in vitro* transcribed that corresponded to human *XIST* repeat E or an RNA corresponding to the anti-sense of the repeat. These were then incubated in Ishikawa nuclear extracts and the ability of human CIZ1 to interact with *XIST* repeat A was assessed by western blot and compared with an anti-sense negative control. Human CIZ1 was specifically detected binding to the sense repeat E, at far greater levels than the antisense control (Figure 5C, Supplementary S8). Similar pulldown experiments with bovine *XIST* repeat E revealed that bovine CIZ1 also interacts with this repeat region (Figure 5D, Supplementary S8).

CIZ1 protein exhibits 79.9% aa identity between human and cow, whilst *XIST* repeat E element only exhibits 54.6% nt identity, therefore the precise nature of the binding could be different between the two species. We hypothesised that given the high level of identity of CIZ1 in the zinc finger domain (Supplementary S9), which is likely to be responsible for RNA binding (Warder and Keherly 2003), human CIZ1 protein would be capable of binding bovine *XIST* repeat E. Therefore, we incubated bovine *XIST* repeat E with human Ishikawa nuclear extract. Low levels of CIZ1 were detected bound to the sense RNA and at similar levels to the anti-sense control (Figure 5E). This suggests that human CIZ1 protein is not able to interact with the bovine E repeat, which is substantially different in sequence.

### Lineage specific positive selection on *XIST* interacting proteins

To further understand the relationship between *XIST* and its protein interactors across placental mammals, we examined whether homologs of the seven protein interacting partners of *XIST* are under similar selective constraints across species. Both site-and lineage-specific models of evolution were employed on a gene-by-gene basis (Yang 2007; Webb, Walsh, and O’Connell 2017) and the human, mouse, cow and pig lineages were labelled as foreground. Following LRT with appropriate null models, the lineages with evidence of positive selection are summarised in **Table 1**, only models of best fit are described below. For CIZ, there was only evidence for positive selection acting in the mouse lineage and not in any of the other three lineages tested (i.e. human, pig and cow). For CIZ it was estimated that ∼60.5% of sites are evolving under strong purifying selection with ω=0.11, and ∼32% of sites are evolving neutrally, and in the mouse lineage alone a further 2.5% of all sites were estimated to evolve under positive selection (ω_2_=4.7). There were 25 amino acid sites with a PP >0.5, two with a PP >0.95, but none above 0.99 (Figure 6). Four out of the 25 amino acids predicted to be under positive selection (702N, 703P, 704S and 713R) spanned the zinc-finger domain of mouse Ciz1 (702-733 aa in mouse Ciz1; Figure 6), and zinc-finger domains have been shown to have RNA binding capacity (Klug 1999; Brown 2005).

**Figure 6:**
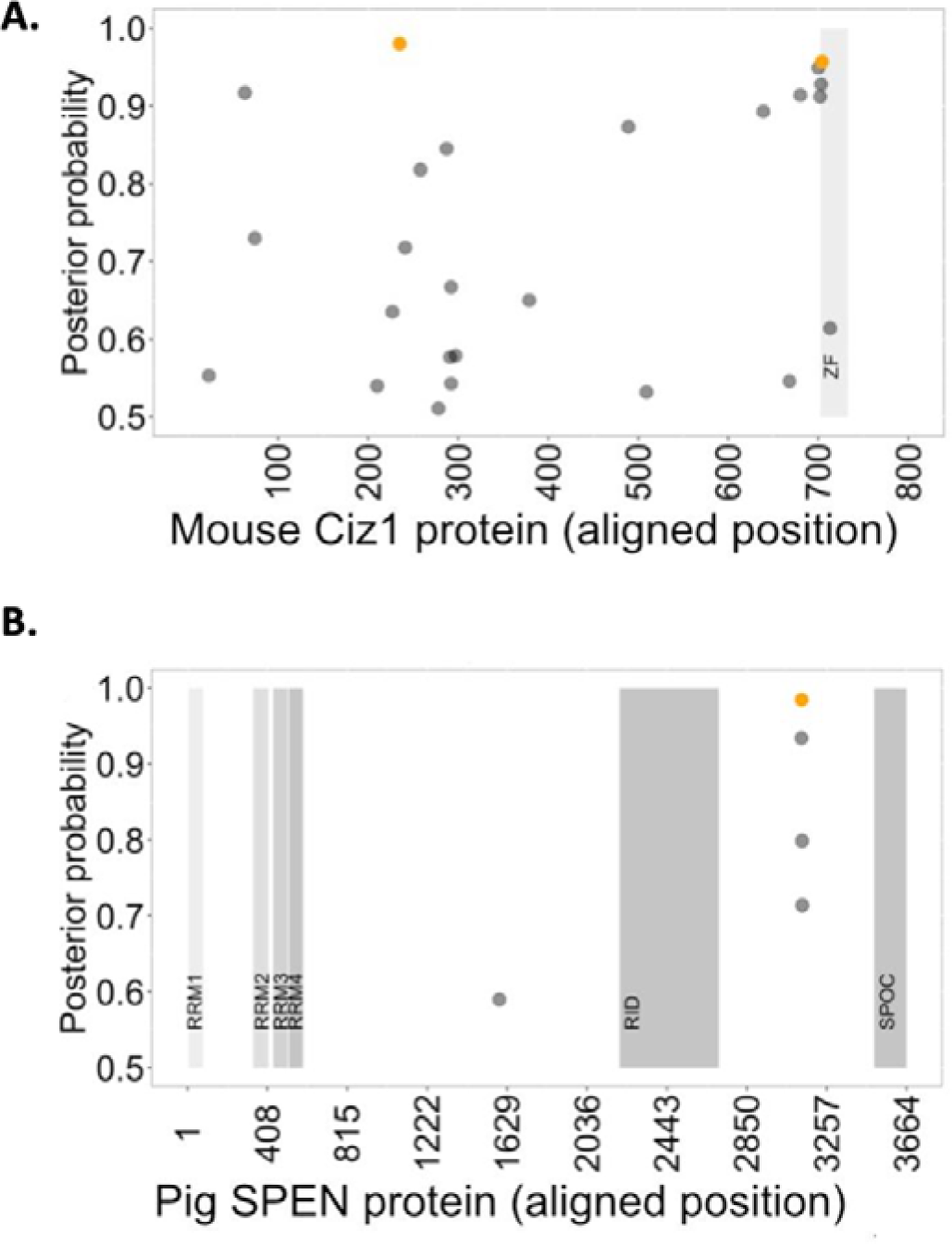
Position of positively selected residues in the proteins encoded by Ciz1 and SPEN. On the *x-axis* is the position in the sequence alignment and on the *y-axis* the posterior probability (BEB) of each position being positively selected (only posterior probabilities ranging >0.5 were considered). The pale grey regions indicate the location of domains of known function. Black dots indicate positively selected sites with a posterior probability score between 0.5 and 0.95, and dots in orange have a posterior probability score between 0.95 and 1.0. **A)** Profile of positively selected residues found only in the mouse lineage for Ciz1. The region highlighted in grey corresponds to the zinc-finger domain (ZF) from position 702 to 733 of the amino acid sequence. **B)** Profile of positively selected residues found only in the pig lineage for SPEN. The vertical grey regions correspond to the known functional domains, left to right these are RNA Recognition Motif (RRM) 1 (6-81aa), RRM2 (335-415aa), RRM3 (438-513aa), RRM4 (517-589), receptor interaction domain (RID) (2201-2707aa) and SPEN paralogue/orthologue C-terminal (SPOC) domain (3498-3664aa).

**Table 1:** Estimates of parameters for lineage specific positive selection.

| Gene:<br>Foreground<br>Lineage | P | Estimates of parameters | Positively<br>selected sites<br>(BEB) |
| --- | --- | --- | --- |
| ***Ciz1*:Mouse** |  |  |  |
| modelA | 7 | p0=0.60542 p1=0.3226 p2=0.04696<br>p3=0.02502 $\omega_0=0.11444$ $\omega_1=1.0$<br>$\omega_2=4.72081$ | 25>0.5,<br>2>0.95, 0>0.99 |
| ***SPEN*: Pig** |  |  |  |
| modelA | 7 | p0=0.80872, p1=0.18868,<br>p2=0.00211, p3=0.00049,<br>$\omega_0=0.08452$, $\omega_1=1.0$, $\omega_2=459.0714$ | 4 >0.5, 3>0.95,<br>2>0.99 |
| ***RBM15*: Cow** |  |  |  |
| modelA | 7 | p0=0.90387 p1=0.09105 p2=0.00462<br>p3=0.00047 $\omega_0=0.03298$ $\omega_1=1.0$<br>$\omega_2=935.93937$ | 3>0.5, 0>0.95,<br>0>0.99 |

The SPEN gene showed evidence of positive selection in the pig lineage with ∼0.2% of sites predicted to be under positive selection (ω_2_=459). There were four sites (1173M, 2711L, 2712Q, 2714Q, 2715Q) with a PP >0.5, none were found to overlap with known functional domains as described by UniProt (RRM1 35-110 aa and RRM2 114-186 aa, Figure 6 - annotated with known human SPEN domains). This would imply that the RNA-binding domains of this protein have been under functional constraint between the species examined here, which is consistent with what others have found (Carter et al. 2020). We also found evidence of positive selection in the RBM15 gene on the cow lineage (Table 1) with ∼0.46% of sites predicted to have ω_2_=935.6 and one of these sites (956F) is within the SPOC domain SPOC, predicting that this domain known to mediate protein-protein interactions potentially has altered function in the cow lineage alone.

## DISCUSSION

*XIST* lncRNA is essential for XCI and is therefore indispensable for proper embryogenesis across placental mammals. The process of XCI has been extensively studied in the mouse model with aspects of the process extrapolated to other placental mammals. However, recent data have demonstrated that mouse development (including X-chromosome inactivation) is sufficiently divergent that it may be considered an outlier in the context of placental mammals (Alberio, Kobayashi, and Surani 2021). Therefore, investigation into whether the protein interacting partners of mouse *Xist* are conserved across mammals is warranted. To determine if the proteins that interact with *XIST* in mouse also interact with *XIST* in other species, we performed detailed *in vitro* experiments using human, pig and bovine as our comparator mammals. We have identified coordinate expression of the protein partners *XIST* interacts with in mouse, in across human, pig and cow by at RNA and protein levels (Figure 2). The majority of data on *XIST* and its partners has been from mouse *in vivo* models, stem cells or more generic human cell lines e.g. HEK293 cells. We used the endometrium as a model as it is more biologically relevant, readily available tissue, and with substantial expression of *XIST* (Figure 2B) increasing the likelihood of being able to pull down proteins that interact with *XIST*. We looked at two species with different embryo morphologies (cow and pig). These species represent different clades from the model systems of mice and humans. They also exhibit different developmental morphology that is more similar to the range of placental mammals than mouse is (Fogarty et al. 2017; Daigneault et al. 2018). The timing and type of XCI is also more comparable to humans, and they undergo a protracted pre-implantation and superficial implantation period of pregnancy (Sahakyan et al. 2017; Petropoulos, et al. 2016; Okamoto, et al. 2011; Yu et al. 2020). These represent more diverse species to investigate whether protein partners of *XIST* are conserved or divergent amongst placental mammals. Even with this diversity of morphology, we have determined that *XIST* and a subset of its predicted protein partners are expressed in the same tissue and therefore are likely to be interacting.

We provide evidence that human *XIST* interacts with the proteins WTAP, SPEN, hnRNPK and RBM15 in human, and that CIZ1 interacts with *XIST* in both humans and bovine via the E-repeat. These data suggest that some of the *XIST* protein partners are conserved in mammals with different with different developmental morphologies, suggesting that the molecular mechanisms may have evolved before the emergence of different developmental pathways in early embryo development and different placental structures.

Technical difficulties with using antibodies generated against human proteins in bovine meant we were unable to determine whether many interactions also occur in cow. RIP is highly sensitive to the quality of antibody used that can bind their target native protein with high affinity within lysate, which is different to an antibody being able to detect a protein in western blot (denatured, partially purified). This may explain why antibodies successfully deployed for westerns from bovine stromal cells (Figure 3C) were not successful at pulling down their antigen.

Here we have provided evidence that CIZ1 interacts with *XIST* through the E repeat in both human and bovine. CIZ1 is very similar between human and cow at the peptide level (85%) but repeat E is quite different at the nucleotide level (55% nt identity). Bovine *XIST* repeat E is also smaller in length than the human E repeat (Yen, et al. 2007). Our results show that human CIZ1 does not bind the bovine repeat E. Given the interaction is conserved across mouse, human and bovine, this is not a species-specific interaction but rather the nature of *XIST*-Ciz1 interaction is species-specific. This could potentially be because of differences in binding kinetics or the number of binding sites with repeat E. An increasing number of CIZ1 binding sites within *XIST* could contribute to more stable binding of *XIST* by CIZ1 (Brockdorff 2019). There are also some amino acid differences within the RNA binding domain of CIZ1, which could provide an explanation for a difference in binding specificity (Supplementary S9). Another potential explanation for the lack of cross-species interaction is that human CIZ1 could require cooperative binding with other, unknown factors, that are not necessary or present in cow. Additionally, the evolution of the *XIST* RNA across placental mammals was coincident with the gain, expansion and exaptation as well as the loss of repetitive elements of various origin (i.e. retroviral or TE elements) across its sequence, which might contribute to differences in motifs within the E repeat between human and cow (Yen et al., 2007, Elisaphenko et al., 2008, Carlevaro-Fita et al., 2019). Given the substantial peri-implantation differences in embryo/conceptus morphology between human and cow it is likely that some of the XIST interactions are not expressed co-ordinately in these species.

The mouse specific signature of positive selection for *CIZ1* is within the known RNA binding zinc finger domain of the protein, suggesting an altered function for Ciz1 unique to the mouse lineage. Specifically, 4 out of 25 aa indicated to be under positive selection are within this zinc-finger domain. The observed functional divergence in mouse may be driven by divergent *Ciz1* function rather than *Xist* repeat E. CIZ1 has an array of other functions outside XCI, which could impact the selective pressures it is under (Nishibe et al. 2013; Pauzaite et al. 2016; Stewart et al. 2019). Interestingly, the repeat E sequence is more similar between human and cow than with mouse. This might likely indicate that mouse *Xist* does not bind to human or bovine repeat E elements. This is also consistent with the observation that the mouse is an outlier in terms of mechanistic output from XCI, with far fewer genes that escape compared with 18 other mammals, including human, cow and pig (Balaton et al. 2021). Recent studies have indicated that differing levels of CIZ1 protein will lead to differences in its ability to generate RNA-protein assemblies in mouse (Sofi et al. 2022). Therefore, differences in the levels of CIZ1 across species and cell types might contribute to differing functional outputs.

We show that regardless of the rapid rates of change we observe in the lncRNA *XIST* across mammals, the underlying interactions between *XIST* and a subset of its protein partners are largely maintained between human and mouse. Indeed, in the case of CIZ1 we report that the conservation of partners is maintained across human, mouse and bovine. However, the precise biochemical basis of this interaction may be different. Despite differences in timing of XCI and early embryo morphologies between mouse, human and cow, CIZ1-*XIST* interaction is maintained. Future work will look to understand how a difference in the nature of CIZ1 binding contributes to the difference in functional output from *XIST*’s role in XCI and how this contributes to timing and type of XCI. Leading on from this work it will be important to understand how differences in timing of XCI linked to differences in embryo morphology are linked to differences in *XIST*-protein complexes.

## Acknowledgements

This work was undertaken on ARC3, part of the High-Performance Computing facilities at the University of Leeds, UK. The authors would also like to thank Peter Mulhair, David T. Orr and Georgios Nikolopoulos for their guidance with the comparative sequence analyses and computational biology aspects of the work. The authors would also like to thank Tayah Hopes, Aikaterini Douka, Karl Norris and Josephina Sampson for advice on molecular biology and Lynne McKeown for mice uterii.

## Author contributions

NF, JLA and MJOC conceived of the study, designed the experiments and interpreted the results. Experiments were designed, performed, analysed and interpreted by IT. HT, IM-E and IT isolated bovine stromal cells from cow endometrial tissue, and isolated mouse and pig uterii. AW provided advice on RIP and RNA pulldowns. IT and MJO’C carried out the comparative and evolutionary sequence analysis and MJO’C interpreted these results. MJO’C, NF and JLA, drafted the manuscript with input from IT, and all authors contributed revising, editing, and critiquing the manuscript.

