## Supplemental Information for "Differences in binding preferences for XIST partners are observed in mammals with different early pregnancy morphologies"

**Supplementary Information**

**Supplementary Table 1 (S1).** List of primers used for RT-qPCR assessment of transcript abundance for XIST and the mRNA of potential protein partners.

| **Species** | **Gene** | **Direction** | **Sequence** |
| --- | --- | --- | --- |
| Human | *XIST* | For | GGCTCCTCTTGGACATTCTGAG |
| Human | *XIST* | Rev | AGCTTGGCCAGATTCTCAAAG |
| Human | *SPEN* | For | GAAGGATGACGGTGGAGACAGA |
| Human | *SPEN* | Rev | CTTGAGGGACTCGGTCTGGC |
| Human | *WTAP* | For | CTAAAGCAACAACAGCAGGAGTC |
| Human | *WTAP* | Rev | GGTACTGGATTTGAGTAGTACACTCT |
| Human | *RBM15* | For | GTCTTCTTGTGGAGGGTTCAACT |
| Human | *RBM15* | Rev | CCCTGCTACTTTGATGCGTC |
| Human | *LBR* | For | AGGAGTACCTGGTGTGTTTCTCAT |
| Human | *LBR* | Rev | CTGGCAAAGGAGGAGGGAA |
| Human | *CIZ1* | For | GAGATGCCAGGGGTATGGG |
| Human | *CIZ1* | Rev | TGGAGGAGACGGAGTCACTGG |
| Human | *hnRNPK* | For | GCGTCCCATGCCTCCATCTAGAAG |
| Human | *hnRNPK* | Rev | CTGAAACCAACCATGCCGTC |
| Human | *hnRNPK* | For | GCTATCCATACCCTCGTGCC |
| Human | *hnRNPK* | Rev | CGTCCTCTGAAGTTCTGGTTGT |
| Human | *ACTB* | For | CTTCCTGGGCATGGAGTCC |
| Human | *ACTB* | Rev | TGATCTTGATCTTCATTGTGCTGG |
| Mouse | *Xist* | For | AGTGGAAATTGGCTGGATTCAG |
| Mouse | *Xist* | Rev | CTTGGTCTTGGGGATAGAAGGA |
| Mouse | *Spen* | For | CTCCAATCAGCCTGCCTACG |
| Mouse | *Spen* | Rev | GTTCAGAGCCTCACACCGAG |
| Mouse | *Wtap* | For | ACCACTCAAATCCAGTACCTCAAG |
| Mouse | *Wtap* | Rev | TTGGGCTTGTTCCAGTTTGTC |
| Mouse | *Rbm15* | For | ACCGATTTGGCACCATTCG |
| Mouse | *Rbm15* | Rev | CCTGAGGCGACGATCTGG |
| Mouse | *Lbr* | For | CAGGAGAGAAGAGGTCAAAGCC |
| Mouse | *Lbr* | Rev | ATGAGGACCGCACCAGGTACT |
| Mouse | *Ciz1* | For | CAAGCAGGTGAAGCCGAG |
| Mouse | *Ciz1* | Rev | TTTGACAGACATAGCCCATCACT |
| Mouse | *hnRNPK* | For | TCCGTACAGACTACAATGCCAG |
| Mouse | *hnRNPK* | Rev | GCCCTCTTCCAAGGTAGG |
| Mouse | *hnRNPU* | For | AGAGGACCGAGTTAGAGGACC |
| Mouse | *hnRNPU* | Rev | CCTGCCACCATCATCTTGTC |
| Mouse | *Actb* | For | GGCACCAGGGTGTGATGG |
| Mouse | *Actb* | Rev | TCCATGTCGTCCCAGTTGG |
| Cow | *XIST* | For | AATCGTTTGTGTTGTGTGAGTGG |
| Cow | *XIST* | Rev | TACTTAGCACAGTTACCCCTCAG |
| Cow | *SPEN* | For | CAGTGACAGCACTGATTCCAGC |
| Cow | *SPEN* | Rev | CGCACTGGAAGATTCTGAACC |
| Cow | *WTAP* | For | GGAACAAGCCCAAAATGAACTG |
| Cow | *WTAP* | Rev | GAGATCAGCAATGGTGGACCC |
| Cow | *RBM15* | For | ACCATACGCACCATTGACTACC |
| Cow | *RBM15* | Rev | GTCTACTCTAAGGCGACGATCTG |
| Cow | *LBR* | For | GCTGGTGCTGAAGCCATTTG |
| Cow | *LBR* | Rev | CCTGTGTGTGTTTGTGAGGCAT |
| Cow | *CIZ1* | For | CACCCGAAGACCAGGAACC |
| Cow | *CIZ1* | Rev | GGCGGCTCAGAGGCTTCA |
| Cow | *hnRNPK* | For | GAATCTTCCTCTTCCACCACC |
| Cow | *hnRNPK* | Rev | CTGAAACCAACCATGCCATCA |
| Cow | *hnRNPU* | For | GGCATTGGCTATCCGTACC |
| Cow | *hnRNPU* | Rev | CGTCCTCTGAAGTTCTGGTTGT |
| Cow | *ACTB* | For | CGCCATGGATGATGATATTGC |
| Cow | *ACTB* | Rev | AAGCCGGCCTTGCACAT |
| Pig | *XIST* | For | AGAAAGGGTGGTGGAATTGGTC |
| Pig | *XIST* | Rev | GGTGCTGACTGGCTGAATAGAG |
| Pig | *SPEN* | For | AGTGACAGAAGAGAAGACCACGG |
| Pig | *SPEN* | Rev | GAGTCCACTTGTTCAGGCTGTTG |
| Pig | *WTAP* | For | CTAGCAACCAAGGAGCAAGAG |
| Pig | *WTAP* | Rev | CGACCATTGTTGATCTCAGTTGG |
| Pig | *RBM15* | For | CAAAGGTGACAGTTGGGCATAC |
| Pig | *RBM15* | Rev | AAGTCTACTCTAAGGCGACGATC |
| Pig | *LBR* | For | GCCTCGGAATGACCTGTC |
| Pig | *LBR* | Rev | TAATGACCACCCAGCCAATCA |
| Pig | *CIZ1* | For | CCAAGACGAGGACCACTTCATC |
| Pig | *CIZ1* | Rev | ATCTCACCTGCTTGCAGAATTCC |
| Pig | *hnRNPK* | For | AAGGAAGCGACTTTGACTGC |
| Pig | *hnRNPK* | Rev | GTCTGAGTGTTCTCCCGAAGTT |
| Pig | *hnRNPU* | For | AACAGAACAGAAAGGCGGAG |
| Pig | *hnRNPU* | Rev | GCGATTTGGCTCTGCTATACT |
| Pig | *ACTB* | For | GCACGGCATCGTCACCAAC |
| Pig | *ACTB* | Rev | GTCCAGACGCAGGATGGC |

**Supplementary Table 2 (S2):** List of antibodies used, with sources and dilutions used.

| **Antibody** | **Protein** | **Species** | **Supplier** | **Dilution** | **Cat. No. #** |
| --- | --- | --- | --- | --- | --- |
| Primary | β-tubulin | Mouse | DSHB | 1:1000-1:5000 | E7-S |
|  | WTAP | Mouse | ProteinTech | 1:1000_ | 60881-1-Ig |
|  | WTAP | Rabbit | Abcam | 1:1000_ | ab195380 |
|  | RBM15 | Rabbit | ProteinTech | 1:1000_ | 10587-1-AP |
|  | hnRNPK | Rabbit | Abcam | 1:1000_ | ab52600 |
|  | hnRNPU | Rabbit | ProteinTech | 1:1000_ | 14599-1-AP |
|  | CIZ1 | Rabbit | ThermoFisher | 1:1000_ | PA5-27625 |
|  | H3K27me3 | Mouse | Abcam | 1:1000_ | ab6002 |
|  | SPEN | Rabbit | Abcam | 1:1000_ | ab72266 |
|  | Lamin B1 | Rabbit | Abcam | 1:1000_ | ab133741 |
| Secondary (HRP-linked) | Mouse IgG | Goat | Cell Signalling  Technology | 1:5000_ | 7076S |
|  | Rabbit IgG | Goat | Cell Signalling  Technology | 1:5000_ | 7074S |

**Supplementary Table 3 (S3): List of primers used for RIP-RT-qPCR.** Information on species, gene, direction and primer sequence are provided.

| Species | Gene and site | Direction | Sequence |
| --- | --- | --- | --- |
| Human | *ACTB* | For | GCACGGCATCGTCACCAAC |
| Human | *ACTB* | Rev | GTCCAGACGCAGGATGGC |
| Human | *U2* | For | GGAGCAGGGAGATGGAATAGG |
| Human | *U2* | Rev | GCACCGTTCCTGGAGGTA |
| Human | *XIST-exon1* | For | GCATAACAGCAGTGGGACTGAC |
| Human | *XIST-exon1* | Rev | AGGTAGTTCACACTATCTAGGAGC |
| Human | *XIST-exon2-3* | For | GGCTCCTCTTGGACATTCTGAG |
| Human | *XIST-exon2-3* | Rev | AGCTTGGCCAGATTCTCAAAG |
| Human | *XIST-exon6* | For | GCTCGGAACTACATGCCC |
| Human | *XIST-exon6* | Rev | ACAGGACTTTATCTCTCTACTCAGC |
| Cow | *ACTB* | For | CGCCATGGATGATGATATTGC |
| Cow | *ACTB* | Rev | AAGCCGGCCTTGCACAT |
| Cow | *RPL19* | For | GGGTATAGGTAAGCGAAAGGG |
| Cow | *RPL19* | Rev | TCACGGTATCGTCTAAGCAGC |
| Cow | *XIST-exon1* | For | CTGCTCTTCTGCGTTGTGG |
| Cow | *XIST-exon1* | Rev | CAGGGATTCCTCTTCTGCC |
| Cow | *XIST-exon5* | For | CCAATCATCATTCTGGACCCTC |
| Cow | *XIST-exon5* | Rev | CTGTCAATTAGCAGGCAGAGC |

**Supplemental Table** **4 (S4)**: Primers used to clone XIST repeats E. All primers target exons and contain restriction enzyme recognition sites (bold) separated with a 3-4 bp ‘stuffer’ sequence (lowercase).

| **Species** | **Amplicon size (nt)** | **Direction** | **Sequence** |
| --- | --- | --- | --- |
| Human | 1881 | For | ***GCGGCCGC****atcatGCACTCTAGCACTTGAGGATAGC* |
| Human |  | Rev | ***GTCGAC****cttgctGAGTAGCGTTGGCACAGTCCA* |
| Bovine | 923 | For | ***GCGGCCGC****taacTGCTCATCACTGTAGTTTGTCTCT* |
| Boving |  | Rev | ***GTCGAC****tcaTGAGTCTTCTATCCAACTCCAGTC* |

**Supplementary Table 5 (S5)**: Generation of biotinylated repeat E RNA fragments from plasmids. Details of restriction enzymes used to linearise plasmids, and RNA polymerases used to generate RNA fragments

| **Species** | **Restriction enzyme** | **Polymerase** | **Orientation** | **size of RNA (nt)** |
| --- | --- | --- | --- | --- |
| Human | EcoRV | SP6 | Sense | 1979 |
| Human | BamHI | T7 | antisense | 1994 |
| Cow | EcoRV | SP6 | Sense | 1036 |
| Cow | BamHI | T7 | antisense | 1021 |

**Supplementary Table 6 (S6)**: **Species included in selective pressure variation analyses.**

| **Common Name** | **Species name** | **Genome Coverage** | **Genome Version (Ensembl v102)** |
| --- | --- | --- | --- |
| Gorilla | *Gorilla gorilla* | x80 | gorGor4 |
| Human | *Homo sapiens* | Deep | GRCh38.p13 |
| Pig | *Sus scrofa* | x65 | Sscrofa11.1 |
| Rabbit | *Oryctolagus cuniculus* | x7.48 | oryCun2 |
| Orangutan | *Pongo pygmaeus* | x6 | PPYG2 |
| Mouse | *Mus musculus* | Deep | GRCm38.p6 |
| Microbat | *Myotis lucifugus* | x7 | myoLuc2 |
| Macaque | *Macaca mulatta* | Deep | MMUL 10 |
| Horse | *Equus caballus* | x88 | Equ Cab 3 |
| Elephant | *Loxodonta africana* | x7 | Loxafr3.0 |
| Dolphin | *Tursiops truncatus* | x2.59 | turTru1 |
| Cow | *Bos taurus* | x80 | ARS-UCD1.2 |
| Dog | *Canis familiaris* | 50x | UMICH_Zoey_3.1 great dane |
| Chimpanzee | *Pan troglodytes* | 55x | Pan_tro_3 |
| Cat | *Felis catus* | x72 | Felis_Catus_9 |
| Armadillo | *Dasypus novemcinctus* | x6 | dasNov3 |
| Platypus | *Ornithorhynchus anatinus* | x58.8 | mOrnAna1.p.v1 |
| Opossum | *Monodelphis domestica* | N/A | ASM229v1 |

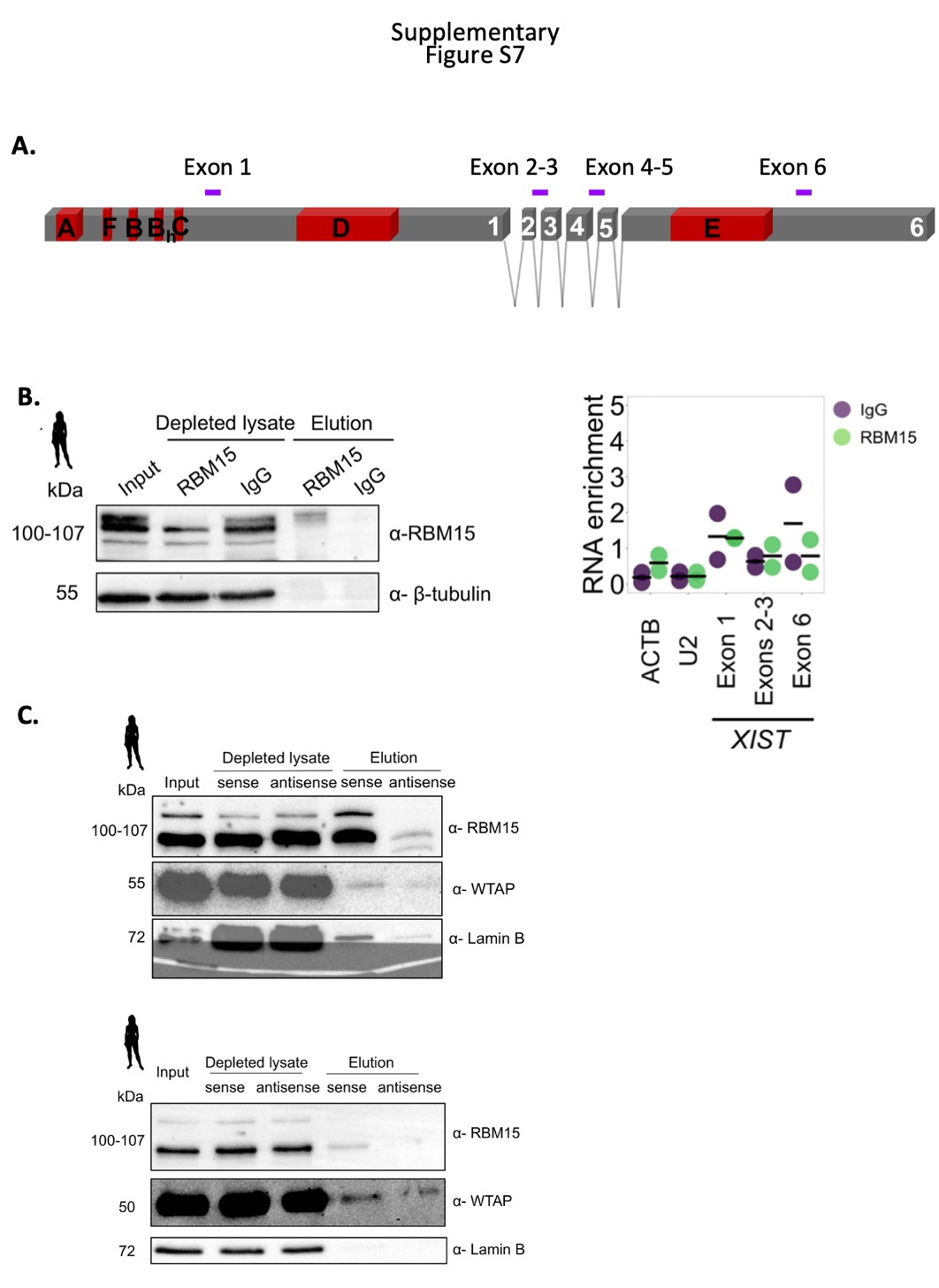

**Supplementary Figure S7:** **A**) Schematic showing the location primers used to detect human *XIST*. Boxes in grey denote exons, with repetitive regions highlighted in red. Numbers on grey boxes denote exon numbers. Amplicons from RT-qPCR are indicated in purple and labelled according to labels in figures. **B**) Representative western blot of RBM15 RIP samples in whole cell extract of ISHIKAWA cells. Bands expected between 100-107 kDa (four isoforms). The amount of protein loaded across input and depleted lysate samples was equivalent to ~500,000 cells and 10% of the elution. n=3 biological replicates. RT-qPCR of RBM15 RIP in whole cell extracts of Ishikawa cells. Fold enrichment of each transcript’s abundance in RIP elutions was normalised to input and reported as RNA enrichment. Three technical replicates were performed for each of two biological replicates. IgG serves as a non-specific negative control in RIP experiments. β-tubulin serves as a non-specific negative control in western blot. *ACTB* and *U2* serve as non-specific transcript negative controls. **C**) *in vitro* transcribed biotinylated human XIST repeat A incubated with nuclear-enriched Ishikawaprotein lysate. Equal amounts of protein were loaded from each sample (~5 μg) and 100% of the elution sample. Input here corresponds to 0.5% of starting amount. Lamin B was used as a negative control. Two independent biological replicates shown.

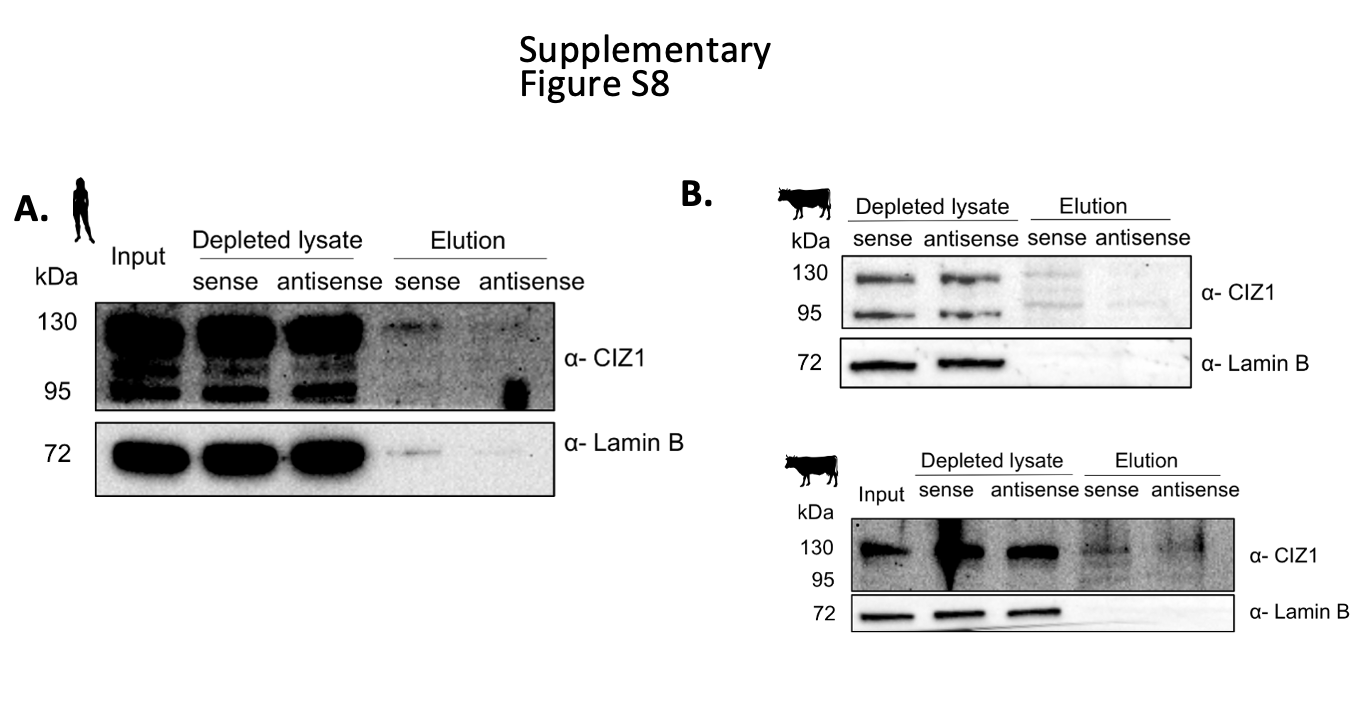

**Supplementary Figure S8:** **A**) Second biological replicate of *in vitro* transcribed biotinylated human *XIST* repeat E incubated with nuclear-enriched Ishikawa protein lysate. Equal amounts of protein were loaded from each sample (~5 μg) and 100% of the elution sample. Input corresponds to 0.5% of starting amount. Lamin B was used as a negative control, n=2 biological replicates. **B**) Second and third biological replicates of *in vitro* transcribed biotinylated bovine *XIST* repeat E incubated with whole cell bovine stromal protein lysates. Equal amounts of protein were loaded from each sample (~5 μg) and 100% of the elution sample. Input here corresponds to 0.5% of starting amount. Lamin B was used as a negative control. Material from different animals was used for each biological replicate.

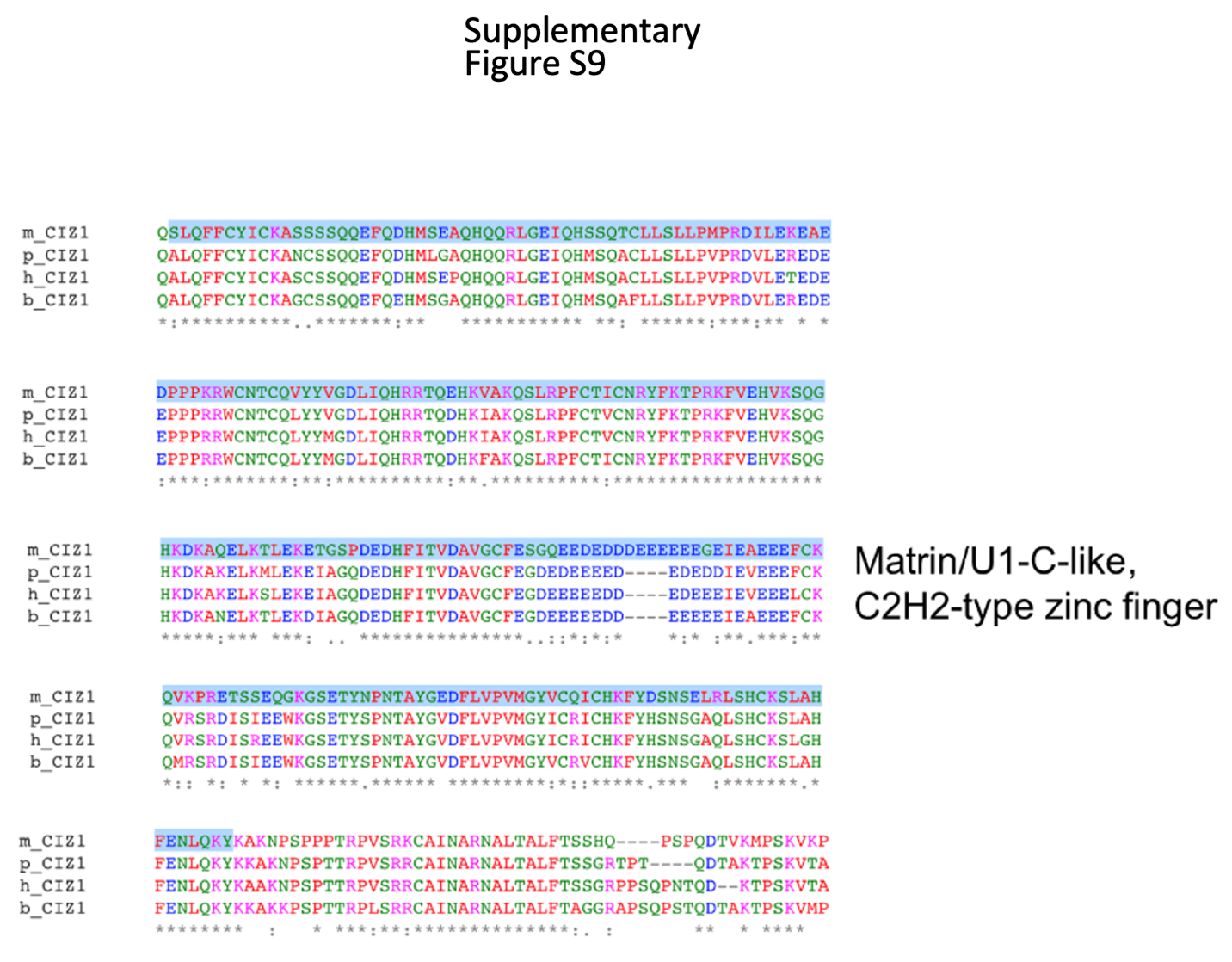

**Supplementary Figure S9:** Alignment of CIZ1 aa sequences from mouse (m), pig (p), human (h) and bovine (b), with the zinger finger domain highlighted in blue.
